# Impact of Advanced Maternal Age on Physiologic Adaptations to Pregnancy in Vervet Monkeys

**DOI:** 10.1101/2020.04.06.027771

**Authors:** Maren Plant, Cecilia Armstrong, Alistaire Ruggiero, Chrissy Sherrill, Beth Uberseder, Rachel Jeffries, Justin Nevarez, Matthew J. Jorgensen, Kylie Kavanagh, Matthew A. Quinn

## Abstract

**Context:** The trend to delay pregnancy in the United States has resulted in the number of advanced maternal age (AMA) pregnancies to also increase. In humans, AMA is associated with a variety of pregnancy-related pathologies such as preeclampsia (PE). While AMA is known to be a factor which contributes to the development of pregnancy-induced diseases, the molecular and cellular mechanisms giving rise to this phenomenon are still very limited. This is due in part to lack of a pre-clinical model which has physiologic relevance to human pregnancy while also allowing control of environmental and genetic variability inherent in human studies.

**Objective:** To determine potential physiologic relevance of the vervet/African green monkey (*Chlorocebus aethiops sabaeus)* as a pre-clinical model to study the effects of AMA on adaptations to pregnancy.

**Design:** Thirteen age-diverse pregnant vervet monkeys (3-16 y.o.) were utilized to measure third trimester blood pressure (BP), complete blood count, iron measurements and hormone levels.

**Results:** Significant associations were observed between third trimester diastolic BP and maternal age. Furthermore, the presence of leukocytosis with enhanced circulating neutrophils was observed in AMA mothers compared to younger mothers. Moreover, we observed a negative relationship between maternal age and estradiol, progesterone and cortisol levels. Finally, offspring born to AMA mothers displayed a postnatal growth retardation phenotype.

**Conclusions:** These studies demonstrate physiologic impairment in the adaptation to pregnancy in AMA vervet/African green monkeys. Our data indicate the vervet/African green monkey may serve as a useful pre-clinical model and tool for deciphering pathological mediators of maternal disease in AMA pregnancy.

## Introduction

Health quality and outcomes for pregnant mothers in the United States are not improving, even with the advancement of modern medicine. In fact, US pregnancy-related maternal mortalities rose 26.6% between 2000 and 2014^1^. Moreover, while the US infant mortality rate is not increasing, it is significantly higher than that of other developed countries^2^. This data highlights a pressing need to understand maternal adaptations to pregnancy in an effort to improve health outcomes for both the mother and child.

Over the last several decades, women and their partners more frequently choose to delay childbirth. The reasons for this change are multi-factorial, but include educational pursuit, access to reliable contraception, and economic uncertainty^3^. While the overall national fertility rate has steadily declined to the lowest numbers recorded in 32 years, the rate of advanced maternal age (AMA) pregnancies, defined as 35 years and older, has risen dramatically^4^. From 2000 to 2014, birth rates for women under 20 declined 42% while the number of women having their first child at age 35 or older rose 23%^5^. The emerging trend of AMA pregnancies is paramount to understand as AMA has been associated with increased risk of several adverse maternal and fetal outcomes^6–9^. For example, AMA is associated with increased risk of gestational diabetes mellitus, placenta previa, and postpartum hemorrhage^7^. In addition, several adverse cardiovascular phenomena have been associated with AMA, including higher risk of developing hypertension and arrhythmias during pregnancy^10^. These conditions are clinically significant considering that 26% of pregnancy-related deaths between 2006 and 2013 had cardiovascular etiologies^10, 11^. Hypertension during pregnancy can also be used to predict future changes for both mother and fetus; women diagnosed with pregnancy-related hypertension experience a 2-8 fold increase in risk for future hypertension,^12–17^while babies born to hypertensive mothers are more likely to develop cardiovascular disease themselves^18–21^. These human data reinforce the need to understand the biological underpinnings of AMA in an effort to improve health outcomes for both mother and child.

Despite the known connection between AMA and pregnancy-related diseases, a gap in knowledge still exists in the pathogenic drivers of this phenomenon in humans. This can somewhat be explained by lack of control over environmental conditions in human studies, along with genetic heterogeneity in human populations. Furthermore, rodent models can lack physiological relevance to reproductive biology in humans. Therefore, a preclinical model with physiological relevance to human pregnancy as well as the ability to control environmental settings is needed to better define underlying mechanisms.

Previous non-human primate (NHP) models have noted similarities between humans and NHPs in hormone physiology during pregnancy and in reproductive biology, which demonstrates their potential as appropriate human pregnancy models^7^. To address this pre-clinical need, we posit and describe herein the use of the vervet/African green monkey (*Chlorocebus aethiops sabaeus)* to model the effects of AMA on maternal adaptation to pregnancy. We demonstrate this model as a pre-clinical platform to garner mechanistic insight, in a tightly controlled environmental setting, into the effects of AMA on pregnancy-induced pathologies, with strong potential for human translational relevance. Our findings demonstrate dysregulated hormonal, cardiovascular, and immunological responses to pregnancy in AMA vervets, all modeling known maladaptive responses to pregnancy in humans. Collectively, our results show that vervets are a clinically relevant model to study the effects of AMA in both maternal and fetal aspects and allow us to compensate for the shortcomings of existing human and animal studies.

## Materials and Methods

### Cohort Selection

A cohort of 13 vervet/African green monkeys (*Chlorocebus aethiops sabaeus)* was selected from the Vervet Research Colony at Wake Forest University School of Medicine. All animals were colony-born, mother-reared, of known-age and were housed in species-typical, matrilineal social groups. Pregnancy status and estimated gestational age was determined via ultrasound as previously described^22^. Modal age of first birth is 4 years old in this colony. Monkeys 3-9 years old were considered optimal maternal age, while monkeys 10 and older were considered to be AMA. In addition, the cohort included primiparous (n=6) and multiparous (n=7) mothers. None of the selected animals exhibited any other comorbidities such as diabetes or heart disease. Other elimination criteria for this study included active participation in other studies. All studies were conducted under the approval of the Institutional Animal Care and Use Committee (IACUC) at Wake Forest School of Medicine.

### Diet

All animals were maintained on a standard chow diet (Monkey Diet Jumbo 5037, LabDiet, St. Louis, MO). Animals were fed *ad libitum* except for fasting on the day of sedated procedures.

### Sedation Protocol

Animals were sedated via intramuscular injections of ketamine (10mg/kg) and midazolam (0.1mg/kg). When necessary, a booster dose (50% of induction dose) was administered to maintain sedation.

### Blood pressure

Systolic and diastolic blood pressure (BP) were measured via high definition oscillometry (S+B medVET, Babenhausen, Germany) as previously desribed^23, 24^. Three high quality measurements were recorded and then averaged to ensure accuracy.

### Complete Blood Counts

Blood was collected via femoral venipuncture into EDTA vacutainers (BD Biosciences; Warwick, RI) approximately two weeks prior to parturition and again 2-5 days postpartum; 500 μL of whole blood were isolated and sent to IDEXX laboratories (Westrbrook, ME) for analysis including a complete blood count (CBC). The remaining blood was centrifuged, and the resulting plasma was collected and stored at −80°C for further analysis.

### Ultrasound

Under sedation, ultrasound (Sonosite M-Turbo; Bothell, WA) was used to measure the biparietal diameter of the fetus *in utero* as previously described^22^. Three measurements were recorded to calculate an average diameter to ensure accuracy.

### Iron Assays

Plasma was analyzed with the BioVision (Milpitas, CA) Total Iron-Binding Capacity (TIBC) and Serum Iron Assay Kit (Colorimetric) according to manufacturer’s instructions. Analysis determined the unbound iron, TIBC + unbound iron, free iron and free iron + transferrin bound iron. These values were used to calculate the TIBC, plasma iron and percent transferrin saturation.

### Hormone Measurements

Plasma was used to determine hormone levels via commercially available enzyme-linked immunosorbent assays for estradiol using the Estradiol Parameter Assay Kit (R&D Systems; Minneapolis, MN, USA) according to manufacturer’s instructions. Progesterone was measured with the Progesterone Human ELISA kit per manufacturer’s protocol (IBL-International; Hamburg, Germany). Finally, cortisol levels were detected utilizing a commercially available kit following manufacturer’s instructions (R&D Systems).

### Statistical Analysis

When comparing two groups an unpaired student’s T-test was used to determine significance. Associations were determined with linear regression analysis. Significance was determined if p<0.05.

## Results

### Maternal Age and Blood Pressure

Given the increased risk for the development of preeclampsia with AMA in humans^25, 26^, we measured BP near the end of the third trimester (approximately two weeks before parturition) in a cohort of age diverse vervets (n=13). Comparing systolic BP with maternal age revealed no significant relationship (R^2^=0.113; p=0.2614) (Fig. 1A). On the other hand, maternal age had a significant positive association with diastolic BP (R^2^=0.3212; p=0.0434) (Fig. 1B). In women, the incidence of preeclampsia decreases substantially in mothers from their first child to their second child^25, 27, 28^. Therefore, we wanted to determine if multiparity might mask the presence of clinical preeclampsia in our AMA cohort. There was a significant positive association between maternal age and number of offspring (*R*^2^=0.9295; p<0.0001) (Supplemental Figure 1). Given the strong association between maternal age and number of offspring we wanted to determine if the protective effects of previous pregnancies are equivalent in young and AMA vervets. This revealed a trend for lower systolic and diastolic BP in young mothers with increasing number of pregnancies (p=0.0554 & p=0.3237 respectively) (Fig. 1C&D). Strikingly, we found in AMA a significant and strong relationship between number of offspring and both diastolic and systolic BP (p=0.0404 & p=0.0014 respectively) (Fig. 1C&D).

**Figure 1:**
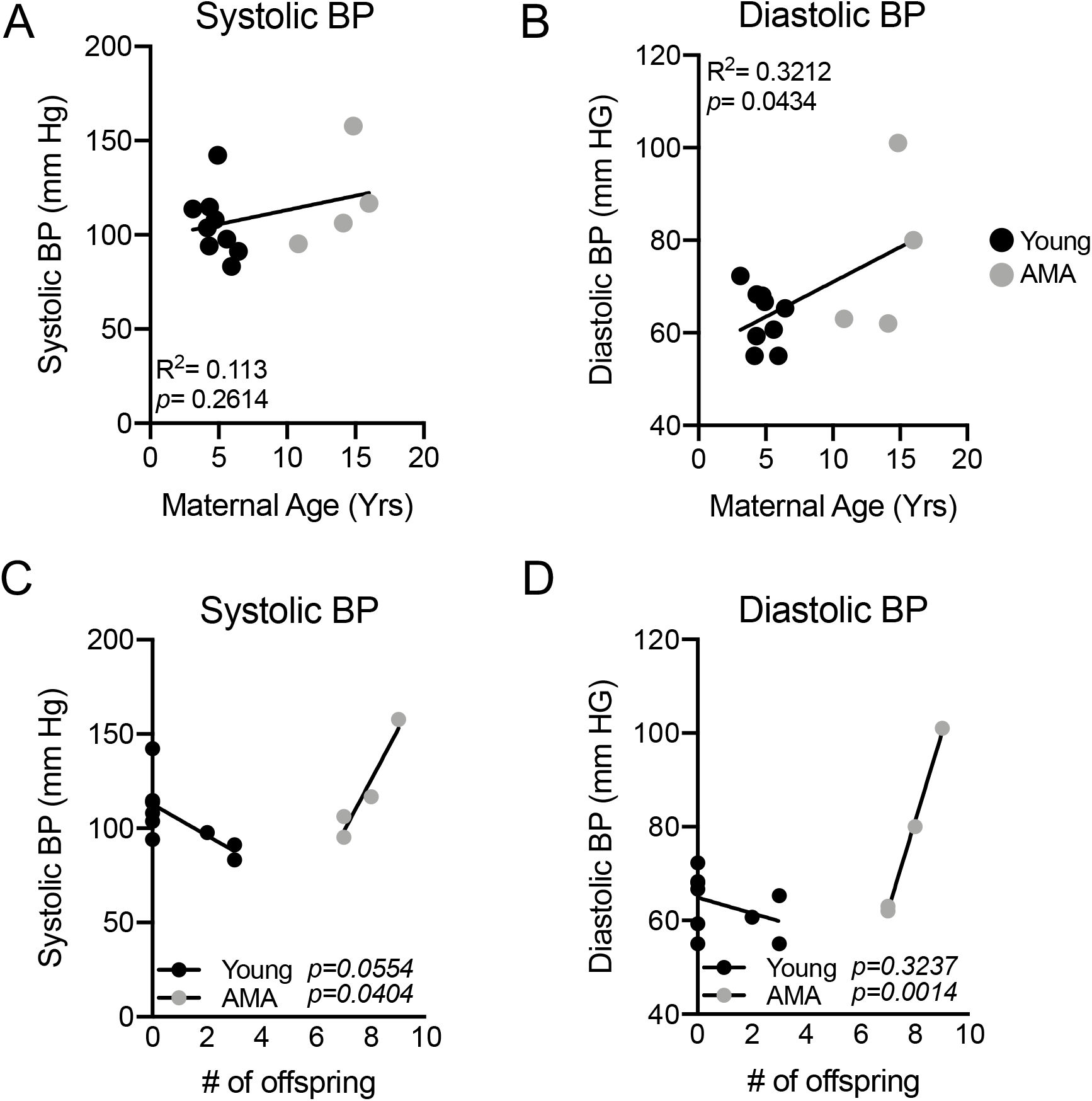
Maternal age is positively associated with third trimester diastolic but not systolic BP. **(A)** Linear regression analysis between third trimester systolic BP and maternal age in vervet monkeys. R^2^=0.1148; *p*=0.2575. **(B)** Linear regression analysis between third trimester diastolic BP and maternal age in vervet monkeys. R^2^=0.3229; *p*=0.0428. **(C)** Linear regression analysis between systolic BP and # of offspring in young (black dots) versus AMA mothers (grey dots). **(D)** Linear regression analysis between diastolic BP and # of offspring in young (black dots) versus AMA mothers (grey dots). N=13 monkeys, 9 young mothers and 4 AMA mothers.

### Leukocytosis in AMA Mothers

Activation of the maternal immune system is a well appreciated contributor to the development of preeclampsia^29–31^. Given the association between maternal age and increasing diastolic BP we sought to determine if maternal age altered third trimester immune cell composition. Complete blood cell counts indicated a significant positive relationship between circulating white blood cell (WBC) number and maternal age (Fig. 2A). Stratifying monkeys between young and AMA revealed significantly higher circulating WBCs in AMA mothers compared to their younger counterparts (Fig. 2B). Our initial screen to determine the cellular components contributing to leukocytosis in AMA mothers revealed no significant alterations in total circulating lymphocyte counts (R^2^=0.02977; p=0.5730) (Supplemental Figure 2).

**Figure 2:**
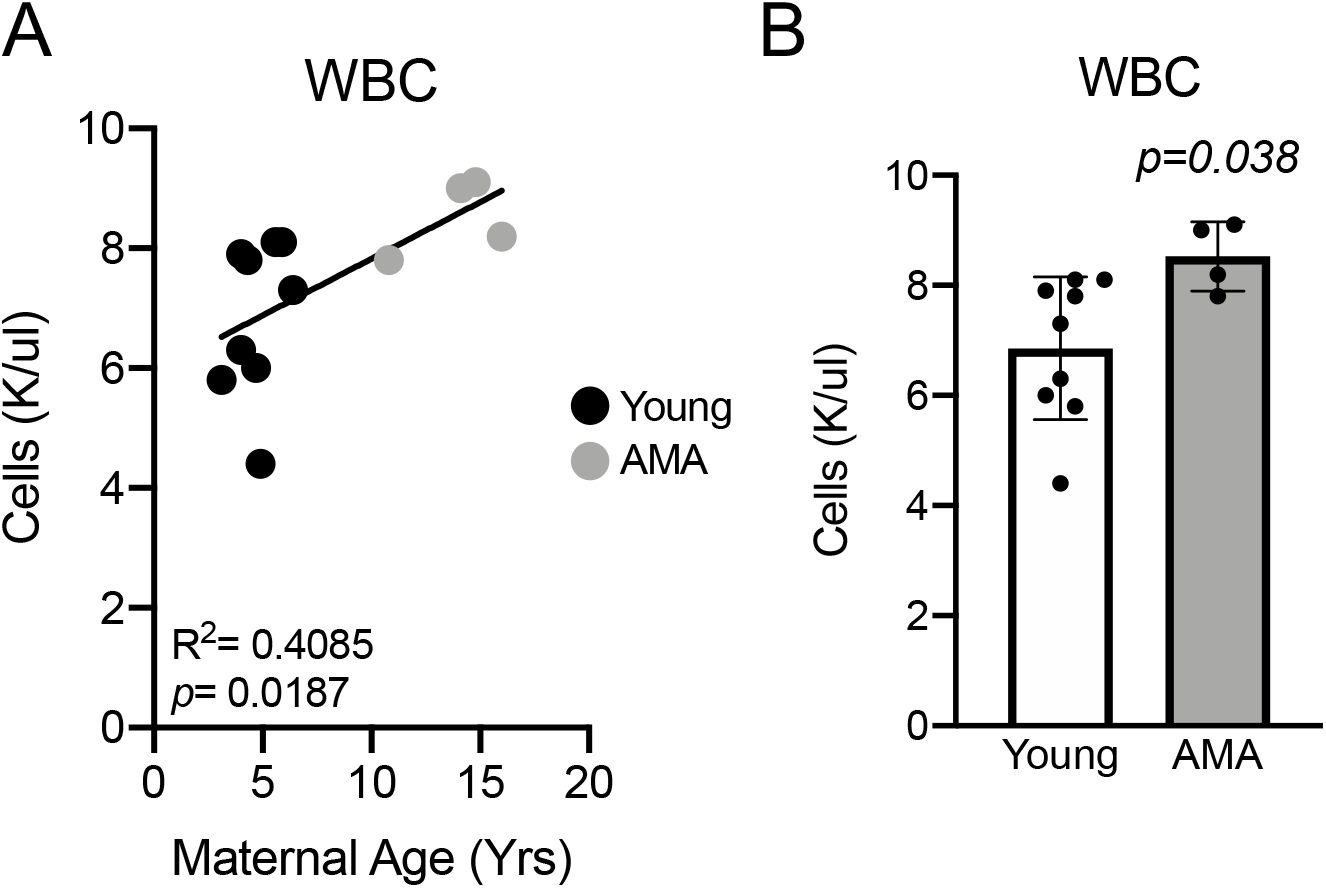
Advanced maternal age promotes third trimester leukocytosis. **(A)** Linear regression of total circulating white blood cell count and maternal age. R^2^=0.4085; *p*=0.0187. N=13 monkeys. **(B)** Third trimester white blood cell count in young (under 10 years of age) and advanced maternal age vervets. N=9 young mothers and 4 advanced maternal age mothers. *p*=*0.038*.

### Stress Leukogram in AMA Mothers

Growing evidence indicates a role for adaptive immune cell activation in the context of preeclampsia^32^. We therefore assessed circulating components of the adaptive immune system including monocytes, basophils, neutrophils and eosinophils. While no alterations were observed in total monocyte (R^2^=0.02997; p=0.6211) and basophil numbers (R^2^=0.01578; p=0.6826) in the circulation related to maternal age (Fig. 3A&B), we observed trends for increased neutrophils with AMA (R^2^=0.2835; p=0.061) (Fig. 3C) and a significant negative association between maternal age and eosinophil numbers (R^2^=0.4016; p=0.02) (Fig. 3D). The presence of neutrophilia and eosinopenia is characteristic of a stress leukogram response^33^.

**Figure3:**
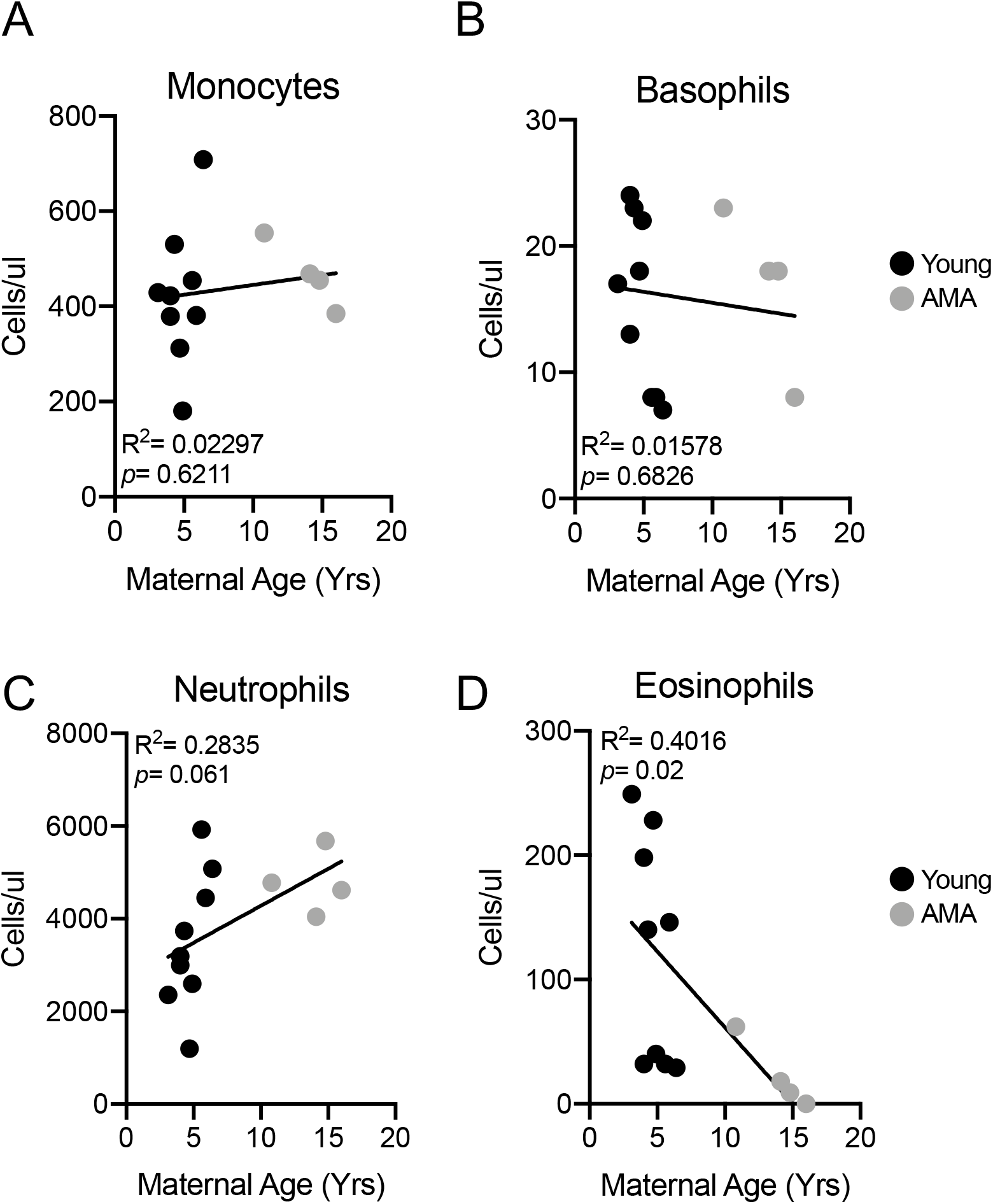
Advanced maternal age is associated with neutrophilia and eosinopenia. **(A)** Linear regression between total circulating monocyte count and maternal age. R^2^=0.02297; *p*=0.6211. **(B)** Linear regression between total circulating basophil count and maternal age. R^2^=0.01578; *p*=0.6826. **(C)** Linear regression between total circulating neutrophil count and maternal age. R^2^=0.2835; *p*=0.061. **(D)** Linear regression between total circulating eosinophil count and maternal age. R^2^=0.4016; *p*=0.02. N=13 monkeys.

### Maternal Body Weight and AMA

To gain insight into mechanisms underlying altered immune and cardiovascular responses we assessed maternal body weight as a risk factor. We observed no significant association between maternal age and maternal pre-pregnancy body weight (Fig. 4).

**Figure 4:**
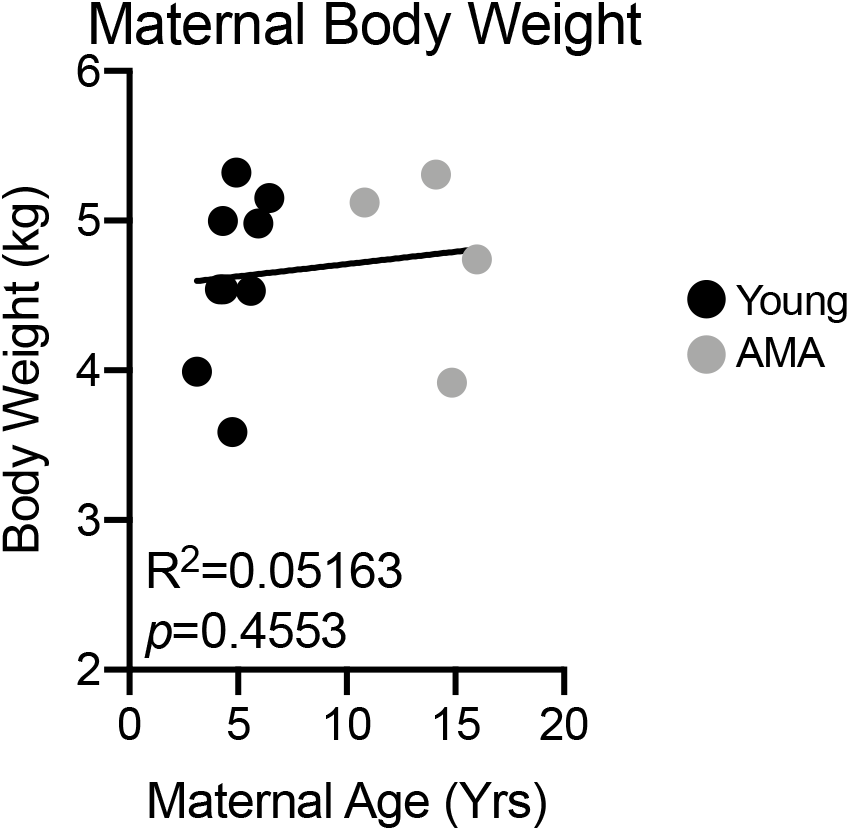
Maternal parity but not body weight is associated with age. **(A)** Linear regression between maternal body weight and maternal age. R^2^=0.05163; *p*=0.4553. N=13 monkeys.

### AMA does not elicit Anemia

We next determined if AMA promotes the development of gestational anemia. We evaluated several parameters associated with anemia in our cohort including red blood cell count, hematocrit and hemoglobin levels. AMA did not alter any biomarker associated with anemia (Fig. 5A-5C). Furthermore, normal serum iron levels (Fig. 5D), total iron binding capacity (Fig. 5E) and % transferrin saturation (Fig. 5F) confirmed the absence of altered iron homeostasis in older mothers.

**Figure 5:**
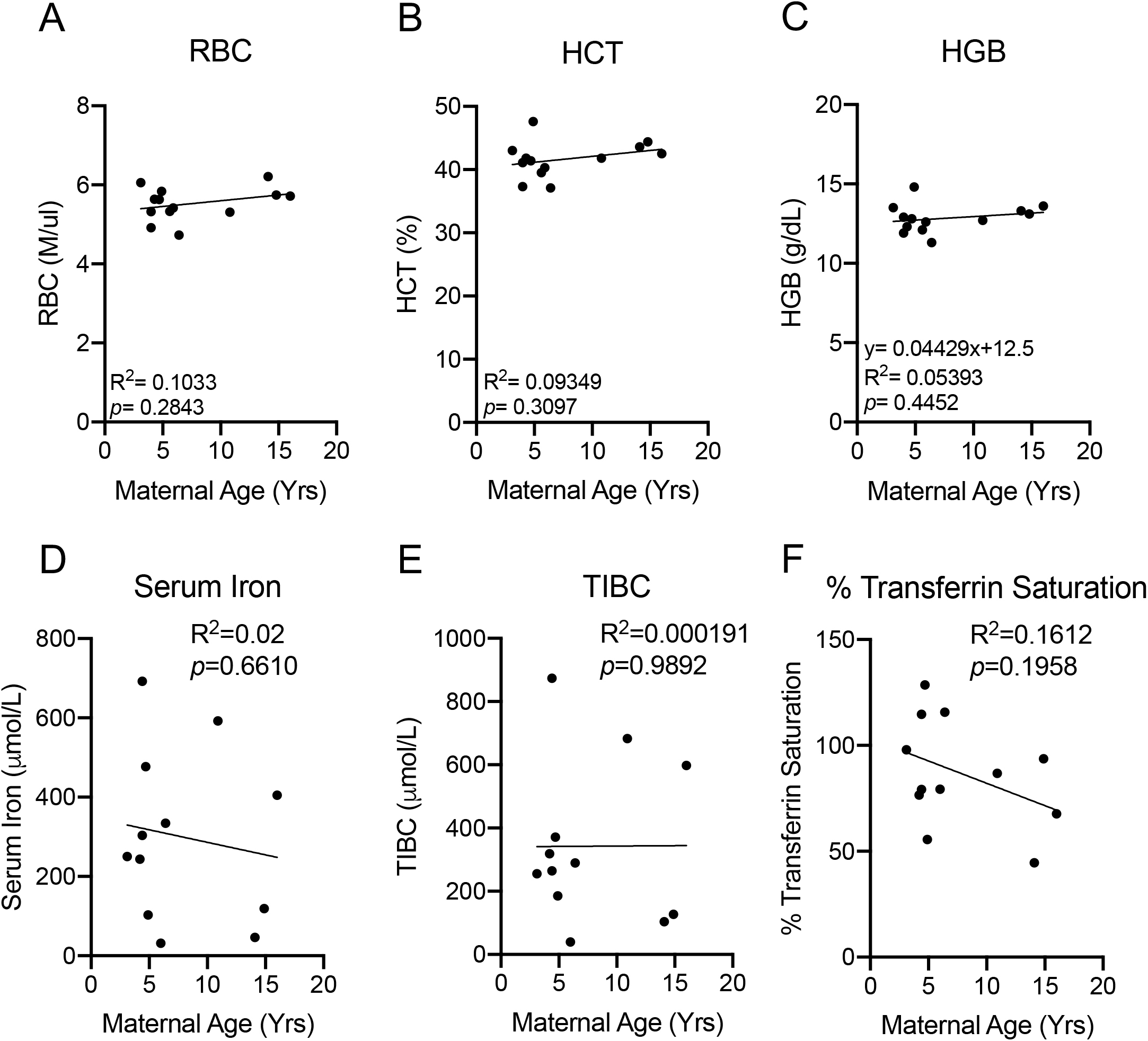
Advanced maternal age does not promote anemia. **(A)** Linear regression between total circulating red blood cell count and maternal age. R^2^=0.1033; *p*=0.2843. **(B)** Linear regression between maternal hematocrit and maternal age. R^2^=0.09349; *p*=0.3097. **(C)** Linear regression between maternal hemoglobin and maternal age. R^2^=0.05393; *p*=0.4452 **(D)** Linear regression between maternal serum iron level and maternal age. R^2^=0.02; *p*=0.6610. **(E)** Linear regression between total iron binding capacity and maternal age. R^2^=0.000191; *p*=0.9892. **(F)** Linear regression between % transferrin saturation and maternal age. R^2^=0.1612; *p*=0.1958. N=13 monkeys.

### Altered Hormonal Responses in AMA Mothers

AMA is associated with low peak gestational estradiol levels^34–36^ and estrogen deficiency has been shown to promote diastolic dysfunction^37^. Therefore, we measured third trimester estradiol levels in our cohort of young and AMA vervets. Enzyme-linked immunosorbent assay (ELISA) revealed AMA mothers had significantly lower third trimester estradiol levels (~60% reduction; R^2^=0.4462; p=0.0176) (Fig. 6A). Further, we found a trend for a negative association between maternal age and circulating third trimester progesterone levels (R^2^=0.2765; p=0.0791) (Fig. 6B). Finally, given the presence of a stress leukogram signature in our AMA mothers, we also measured cortisol levels, revealing a significant negative relationship (R^2^=0.5832; p=0.0038)) between maternal age and third trimester cortisol levels (Fig. 6C).

**Figure 6:**
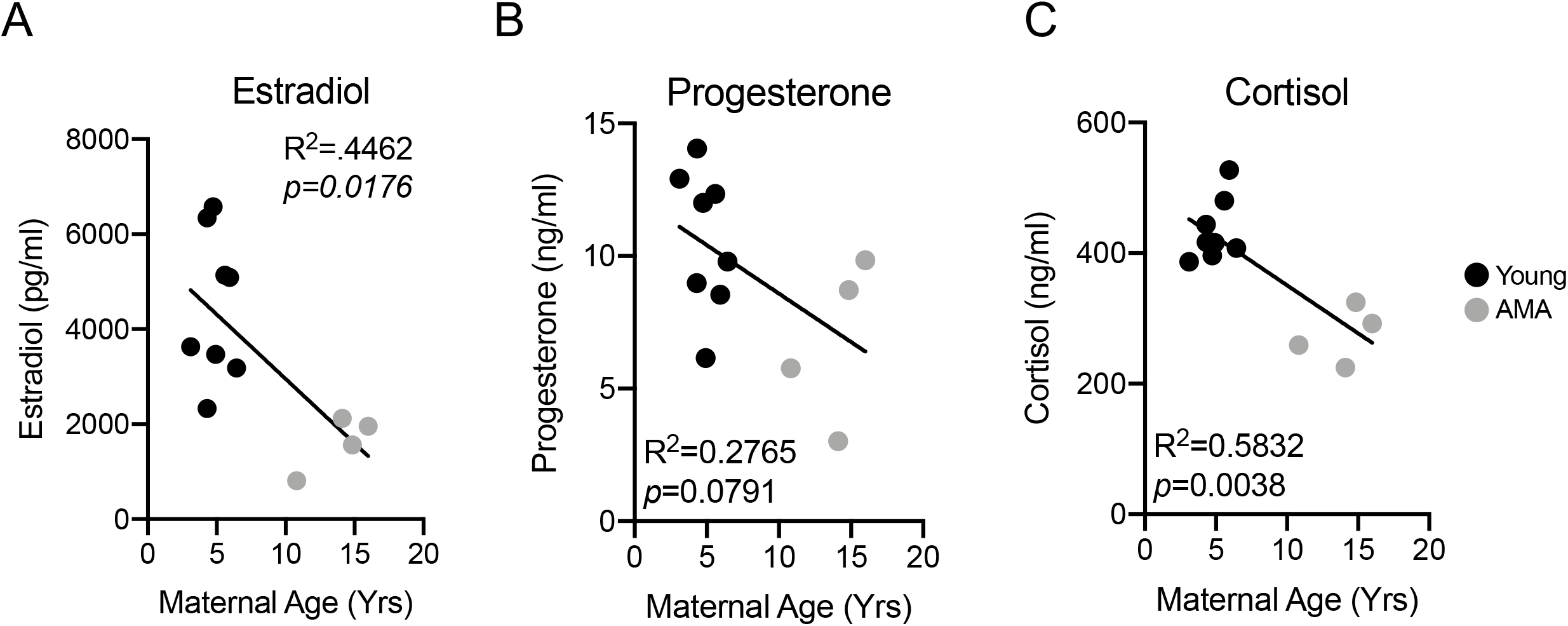
Advanced maternal age disrupts hormonal responses to pregnancy. **(A)** Linear regression between estradiol and maternal age. **(B)** Linear regression between progesterone and maternal age. **(C)** Linear regression between cortisol and maternal age. N=13 monkeys.

### Postnatal Growth Retardation in Offspring from AMA Mothers

We measured fetal biparietal diameter approximately two weeks prior to delivery via ultrasound. No appreciable differences were observed in fetal biparietal diameter within our cohort (Fig. 7A). Accordingly, we also did not observe significant differences in infant body weights between young and AMA age mothers at birth (Fig. 7B). However, following archival growth trajectories over approximately the first year of life in a separate cohort of animals (n=28 young and n=14 aged) revealed significant growth retardation in infants born to AMA mothers (Fig. 7B).

**Figure 7:**
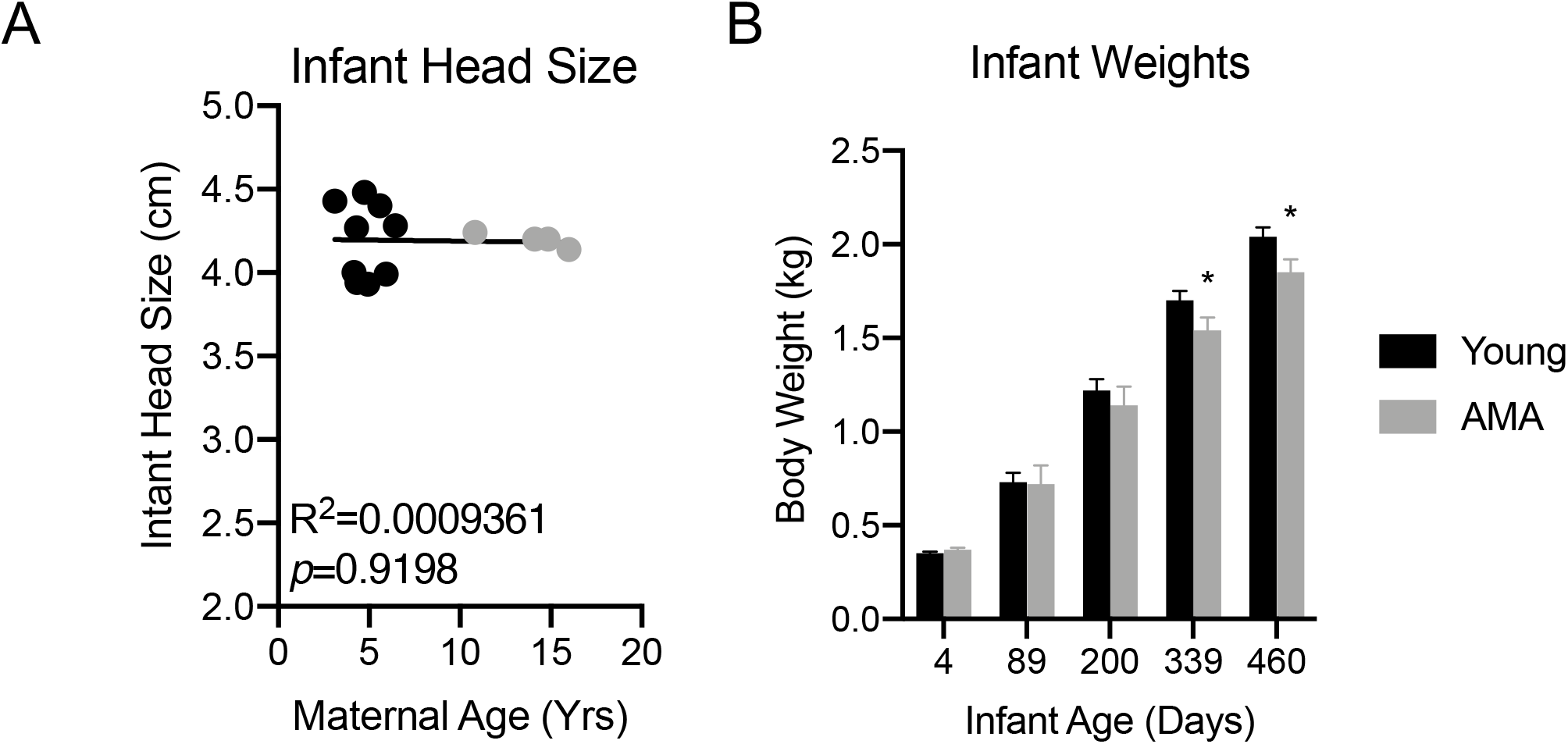
Offspring from advanced maternal age vervets present postnatal growth retardation. **(A)** Linear regression between fetal biparietal diameter and maternal age. R^2^=0.03142; *p*=0.05624. N=13 monkeys. **(B)** Archival growth rates of offspring from a separate cohort of young and advanced maternal age mothers in an expanded cohort of Vervet monkeys. n=28 young and n=14 aged. * denotes *p*<0.05.

## Discussion

AMA in humans is an established risk factor for the development of an array of pregnancy-induced pathologies^6, 8, 26, 38, 39^. While the relationship between maternal age and the incidence of pregnancy-related pathologies exists, pre-clinical models with similar reproductive physiology to that of humans are severely lacking. The current study clearly shows that AMA is associated with disruptions in physiological adaptations to pregnancy in vervet/African green monkeys. In particular, we found the cardiovascular system, immune system and endocrine system all display deficits in responses to pregnancy, suggesting the presence of maternal pathologies in older vervet monkeys. Additionally, first year growth trajectories were impaired in infants born to AMA mothers. These data collectively indicate the vervet monkey as a physiologically relevant pre-clinical model to study the effects of AMA on both maternal and offspring outcomes.

Human studies have revealed a selective increase in third trimester diastolic blood pressure and a decrease in systolic BP with increased maternal age^40^. Consistent with these findings, we observed maternal age to be significantly positively associated with diastolic BP in our vervet model. Contrary to the human studies however, we observed no relationship between age and third trimester systolic BP. These findings indicate that the vervet monkey recapitulates some, but not all aspects of altered BP regulation during pregnancy in older mothers. Gaillard *et al.* indicated that a woman’s maternal age per se was not consistently correlated with gestational hypertension, and that maternal body mass index might influence alterations in BP regulation during pregnancy^38^. In fact, maternal obesity has been shown to interact with maternal age to promote a variety of other pregnancy-induced pathologies^38^. We observed no association between maternal body weight and maternal age (Figure 4), which may explain differences observed between our study in vervet monkeys and human studies in the regulation of third trimester systolic BP.

Beyond elevated BP, a significant immunological component to preeclampsia exists^30–32, 41–43^. While leukocytosis occurs during normal pregnancy^44^, exaggerated leukocytosis occurs in preeclamptic patients^45^. Our observation in the vervet monkey that AMA mothers have significantly elevated white blood cell counts coupled to the presence of diastolic hypertension are consistent with hallmarks of human preeclampsia. Intriguingly, leukocytosis present in humans with preeclampsia is due to an increase in circulating neutrophils counts^45^. Similar to our other data supporting physiological relevance of vervet monkeys to humans for studying the effects of AMA, the older mothers exhibited a higher degree of neutrophilia present in their third trimester compared to young mothers, potentially exacerbating a state of mild preeclampsia.

We did observe a significant positive association in our cohort between maternal age and parity. The elevated parity in our AMA could actually be providing a protective mechanism against the development of more severe preeclampsia, as this disease is more prevalent amongst primiparous mothers^25, 27, 28^. We observed an uncoupling of number of previous offspring and blood pressures between young and AMA mothers. Our data suggest that previous pregnancies are associated with lowered blood pressures in younger mothers; however, in AMA mothers the number of pregnancies was positively associated with both diastolic and systolic BP. These data suggest that either AMA disrupts the protective mechanisms afforded by previous pregnancies, or, that after a certain threshold of previous pregnancies the protective mechanism of parity is lost. Parity has also been associated with immunological tolerance to certain infections during pregnancy such as malaria^46–48^ and multiparity has been demonstrated to confer immunotolerance in rodent models of stroke^49^, indicating a protective role to maternal health in multiparous mothers. While not tested in the current study, further investigation into AMA primiparous third trimester physiology is warranted to determine if multiparity is protective against the development of clinical preeclampsia.

Another known risk factor for the development of preeclampsia in humans is the presence of pregnancy-induced anemia^50–52^. Furthermore, maternal age and parity have been shown to be associated with the presence of anemia in humans^53, 54^. However, we did not observe such associations between anemia and maternal age and multiparity in our study. One explanation for the lack of association between maternal age and anemia in our study is due to diet; while maternal age is associated with the development of anemia in humans, this is largely due to insufficient iron intake during pregnancy^55–57^. Our vervet diet has high levels of iron (230 ppm), which could potentially compensate for AMA as a risk factor.

Estradiol is a well-known cardioprotective hormone. In the non-pregnant state, low estradiol levels, such as those observed during menopause, promote the development of cardiovascular disease^58, 59^. Specifically, postmenopausal women are the primary clinical population diagnosed with heart failure with preserved ejection fraction (HfpEF)^60–62^. The cardioprotective effects of estradiol in preventing HfpEF in estrogen deficient females has been extended to nonhuman primates such as cynomolgus macaques^37^. In the pregnant state, low estrogen levels have been associated with preeclampsia in humans^63–67^. We found AMA is associated with third trimester estrogen deficiency in vervet monkeys, consistent with human data indicating maternal age is negatively correlated with low peak estradiol levels ^34–36^. At the molecular level, estrogens have been shown to antagonize the effect of stress hormones^68–71^. We have demonstrated previously that the antagonistic nature of estrogen on stress hormones is essential for appropriate adaptations to pregnancy and proper fetal development in rodents^68^. Our data indicate AMA disrupts the cortisol/estradiol axis through impaired estradiol production. Furthermore, the presence of a stress leukogram in AMA vervets is suggestive of aberrant stress hormone signaling in aged pregnant vervets^33^.

Maternal stress in humans, like AMA, underlies long-term predisposition of offspring to disease into adulthood. This concept is known as the developmental origin of disease^72^. A commonality between maternal stress and AMA is they are both risk factors for the development of intrauterine growth restriction in humans and small gestational age infants^6, 8, 38, 39, 73, 74^. Our ultrasound data of fetal biparietal diameter revealed no association between maternal age and head size. Furthermore, infant weight at four days post-delivery was comparable between young and AMA mothers. In humans, one driver of the small gestational phenotype is pre-term delivery^75–78^. This may be a possible explanation for why we did not observe low birth weights in vervets, since AMA did not elicit pre-term delivery in our cohort. Beyond low birth weights, prenatal maternal stress in humans dramatically alters postnatal growth rates of offspring. Intriguingly, the offspring growth rate phenotype is dictated by timing of maternal stress, with early stress typically leading to increased growth rates and late stress promoting decreased growth rates in offspring across 21 different mammalian species^79^. Our results of normal infant weight but blunted postnatal growth is suggestive that AMA in vervets corroborates human data resultant of a maternal stress response late during gestation. An additional factor within the paradigm of maternal stress is maternal investment during lactation^79^. We did not cross foster or perform behavioral analyses in our young and AMA vervets post-delivery, therefore we cannot determine if AMA alters maternal investment during the nursing period.

Human studies limit the ability to establish disease causality. Rodent studies on the other hand allow for experimental manipulation to test mechanisms underlying disease, but their reproductive physiology is dramatically different than that of humans. Utilizing an experimental model with direct physiological relevance would allow circumvention of these hurdles. Establishing the vervet monkey as a physiologically relevant preclinical model allows for the ability to tightly regulate environmental conditions and to collect longitudinal measurements, tissues and cells currently not feasible in human studies. This model will allow for the mechanistic dissection of how maternal age promotes pregnancy-induced pathologies with high likelihood for clinical translation and the ability to impact human health.

One primary strength of our study is the establishment of a pre-clinical model with reproductive physiologic relevance to humans for studying the effects of aging on maternal health outcomes. Furthermore, the utilization of clinically relevant assays to characterize the impact of maternal age on adaptations to pregnancy is another primary strength of our study. One weakness with our study is that we focused only on third trimester physiology. It is of the utmost importance to further delineate the effects of AMA during gestation. Moreover, our studies are observational and descriptive in nature. Future studies assessing the effects of estrogen supplementation in AMA vervets on amelioration of cardiac and immunological responses to pregnancy are much needed. Finally, the study may not be powered for certain comparisons, leading to a Type II error such as maternal body weight and anemia related factors.

Our data demonstrate that AMA in vervets summarizes several maladaptive responses observed in humans, particularly dysregulation of hormonal, cardiovascular and immunological responses to pregnancy, and establishes this model for further elucidation of the mechanisms involved in the stress responses involved in maternal adaptation to pregnancy and postnatal growth retardation in humans.

## Acknowledgements

The authors would like to thank M. Christina May Long and Justin Herr for assistance with the Vervet Research Colony, and Dr. Tom Register and Ms. Maryanne Post for their technical assistance with estradiol measurements. We would also like to thank the Biomarker Analytical Core of Wake Forest University Health Sciences for their assistance with cortisol and progesterone measurements.

---

**Supplemental Figure 1:**
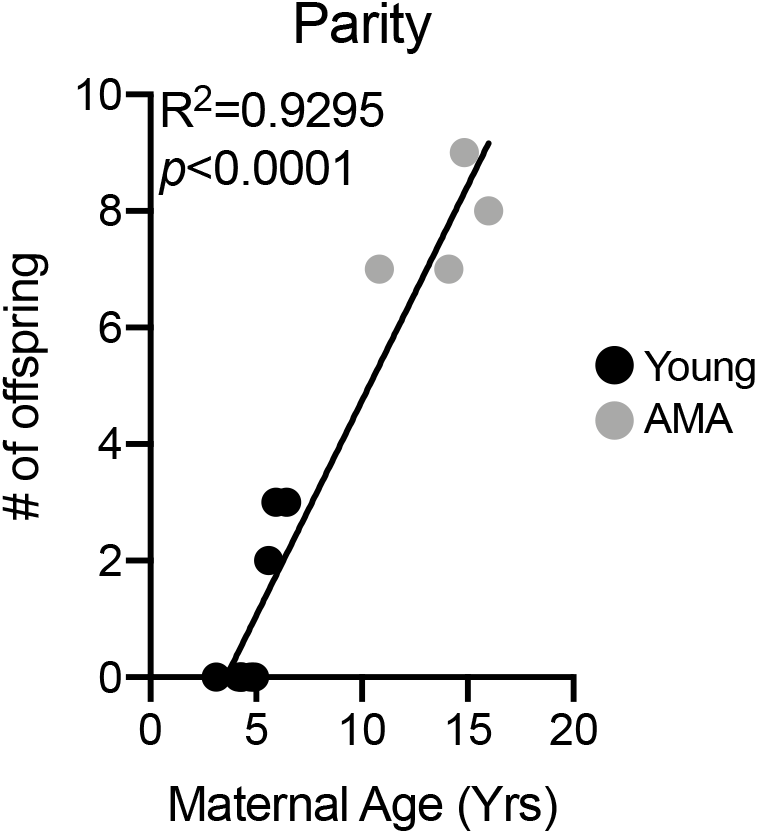
Significant association between maternal age and number of offspring in studied cohort. Linear regression between parity and maternal age. R^2^=0.935; *p*<0.0001. N=13 monkeys.

**Supplemental Figure 2:**
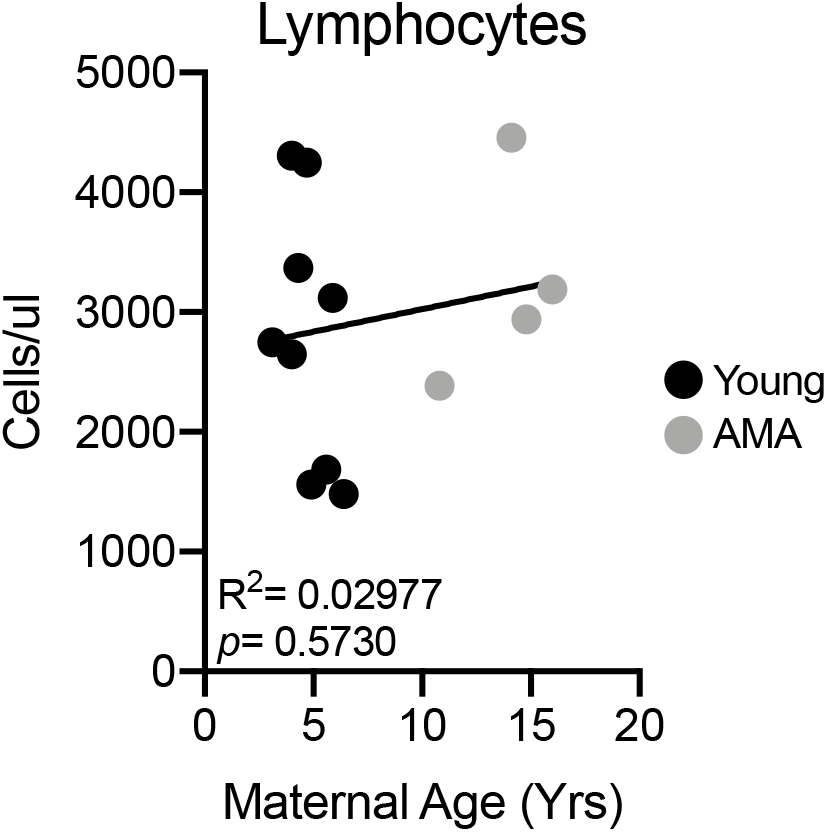
Maternal age does not alter circulating lymphocyte counts. Linear regression between total lymphocyte count and maternal age. R^2^=0.02977; *p*=0.5730. N=13 monkeys.

## References

[1] MacDorman MF, Declercq E, Cabral H, Morton C: Recent Increases in the U.S. Maternal Mortality Rate: Disentangling Trends From Measurement Issues. Obstet Gynecol 2016, 128:447–55.

[2] He X, Akil L, Aker WG, Hwang HM, Ahmad HA: Trends in infant mortality in United States: a brief study of the Southeastern states from 2005-2009. Int J Environ Res Public Health 2015, 12:4908–20.

[3] Mills M, Rindfuss RR, McDonald P, te Velde E, Reproduction E, Society Task F: Why do people postpone parenthood? Reasons and social policy incentives. Hum Reprod Update 2011, 17:848–60.

[4] Brady E. Hamilton JAM, Michelle Osterman, Lauren Rossen: Births: Provisional Data for 2018. Vital Statistics Rapid Release 2019, 7.

[5] Hamilton TJMaBE: Mean Age of Mothers is on the Rise: United States, 2000-2014. NCHS Data Brief 2016, 232.

[6] Lean SC, Derricott H, Jones RL, Heazell AEP: Advanced maternal age and adverse pregnancy outcomes: A systematic review and meta-analysis. PLoS One 2017, 12:e0186287.

[7] Yogev Y, Melamed N, Bardin R, Tenenbaum-Gavish K, Ben-Shitrit G, Ben-Haroush A: Pregnancy outcome at extremely advanced maternal age. Am J Obstet Gynecol 2010, 203:558 e1–7.

[8] Khalil A, Syngelaki A, Maiz N, Zinevich Y, Nicolaides KH: Maternal age and adverse pregnancy outcome: a cohort study. Ultrasound Obstet Gynecol 2013, 42:634–43.

[9] Carolan MC, Davey MA, Biro M, Kealy M: Very advanced maternal age and morbidity in Victoria, Australia: a population based study. BMC Pregnancy Childbirth 2013, 13:80.

[10] De Viti D, Malvasi A, Busardo F, Beck R, Zaami S, Marinelli E: Cardiovascular Outcomes in Advanced Maternal Age Delivering Women. Clinical Review and MedicoLegal Issues. Medicina (Kaunas) 2019, 55.

[11] James AH, Jamison MG, Biswas MS, Brancazio LR, Swamy GK, Myers ER: Acute myocardial infarction in pregnancy: a United States population-based study. Circulation 2006, 113:1564–71.

[12] Behrens I, Basit S, Melbye M, Lykke JA, Wohlfahrt J, Bundgaard H, Thilaganathan B, Boyd HA: Risk of post-pregnancy hypertension in women with a history of hypertensive disorders of pregnancy: nationwide cohort study. BMJ 2017, 358:j3078.

[13] Heida KY, Franx A, van Rijn BB, Eijkemans MJ, Boer JM, Verschuren MW, Oudijk MA, Bots ML, van der Schouw YT: Earlier Age of Onset of Chronic Hypertension and Type 2 Diabetes Mellitus After a Hypertensive Disorder of Pregnancy or Gestational Diabetes Mellitus. Hypertension 2015, 66:1116–22.

[14] Timpka S, Stuart JJ, Tanz LJ, Rimm EB, Franks PW, Rich-Edwards JW: Lifestyle in progression from hypertensive disorders of pregnancy to chronic hypertension in Nurses’ Health Study II: observational cohort study. BMJ 2017, 358:j3024.

[15] Bokslag A, Teunissen PW, Franssen C, van Kesteren F, Kamp O, Ganzevoort W, Paulus WJ, de Groot CJM: Effect of early-onset preeclampsia on cardiovascular risk in the fifth decade of life. Am J Obstet Gynecol 2017, 216:523 e1–e7.

[16] Benschop L, Duvekot JJ, Versmissen J, van Broekhoven V, Steegers EAP, Roeters van Lennep JE: Blood Pressure Profile 1 Year After Severe Preeclampsia. Hypertension 2018, 71:491–8.

[17] Brouwers L, van der Meiden-van Roest AJ, Savelkoul C, Vogelvang TE, Lely AT, Franx A, van Rijn BB: Recurrence of pre-eclampsia and the risk of future hypertension and cardiovascular disease: a systematic review and meta-analysis. BJOG 2018, 125:1642–54.

[18] Kajantie E, Eriksson JG, Osmond C, Thornburg K, Barker DJ: Pre-eclampsia is associated with increased risk of stroke in the adult offspring: the Helsinki birth cohort study. Stroke 2009, 40:1176–80.

[19] Davis EF, Lazdam M, Lewandowski AJ, Worton SA, Kelly B, Kenworthy Y, Adwani S, Wilkinson AR, McCormick K, Sargent I, Redman C, Leeson P: Cardiovascular risk factors in children and young adults born to preeclamptic pregnancies: a systematic review. Pediatrics 2012, 129:e1552–61.

[20] Jayet PY, Rimoldi SF, Stuber T, Salmon CS, Hutter D, Rexhaj E, Thalmann S, Schwab M, Turini P, Sartori-Cucchia C, Nicod P, Villena M, Allemann Y, Scherrer U, Sartori C: Pulmonary and systemic vascular dysfunction in young offspring of mothers with preeclampsia. Circulation 2010, 122:488–94.

[21] Tripathi RR, Rifas-Shiman SL, Hawley N, Hivert MF, Oken E: Hypertensive Disorders of Pregnancy and Offspring Cardiometabolic Health at Midchildhood: Project Viva Findings. J Am Heart Assoc 2018, 7.

[22] Kavanagh K, Dozier BL, Chavanne TJ, Fairbanks LA, Jorgensen MJ, Kaplan JR: Fetal and maternal factors associated with infant mortality in vervet monkeys. J Med Primatol 2011, 40:27–36.

[23] Strawn WB, Chappell MC, Dean RH, Kivlighn S, Ferrario CM: Inhibition of early atherogenesis by losartan in monkeys with diet-induced hypercholesterolemia. Circulation 2000, 101:1586–93.

[24] Strawn WB, Ferrario CM: Angiotensin II AT1 receptor blockade normalizes CD11b+ monocyte production in bone marrow of hypercholesterolemic monkeys. Atherosclerosis 2008, 196:624–32.

[25] Duckitt K, Harrington D: Risk factors for pre-eclampsia at antenatal booking: systematic review of controlled studies. BMJ 2005, 330:565.

[26] Bianco A, Stone J, Lynch L, Lapinski R, Berkowitz G, Berkowitz RL: Pregnancy outcome at age 40 and older. Obstet Gynecol 1996, 87:917–22.

[27] Luo ZC, An N, Xu HR, Larante A, Audibert F, Fraser WD: The effects and mechanisms of primiparity on the risk of pre-eclampsia: a systematic review. Paediatr Perinat Epidemiol 2007, 21 Suppl 1:36–45.

[28] Saftlas AF, Beydoun H, Triche E: Immunogenetic determinants of preeclampsia and related pregnancy disorders: a systematic review. Obstet Gynecol 2005, 106:162–72.

[29] Chatterjee P, Weaver LE, Chiasson VL, Young KJ, Mitchell BM: Do double-stranded RNA receptors play a role in preeclampsia? Placenta 2011, 32:201–5.

[30] Redman CW, Sacks GP, Sargent IL: Preeclampsia: an excessive maternal inflammatory response to pregnancy. Am J Obstet Gynecol 1999, 180:499–506.

[31] Saito S, Shiozaki A, Nakashima A, Sakai M, Sasaki Y: The role of the immune system in preeclampsia. Mol Aspects Med 2007, 28:192–209.

[32] Boucas AP, de Souza BM, Bauer AC, Crispim D: Role of Innate Immunity in Preeclampsia: A Systematic Review. Reprod Sci 2017, 24:1362–70.

[33] Latimer KS, Rakich PM: Clinical interpretation of leukocyte responses. Vet Clin North Am Small Anim Pract 1989, 19:637–68.

[34] Sharma V, Riddle A, Mason BA, Pampiglione J, Campbell S: An analysis of factors influencing the establishment of a clinical pregnancy in an ultrasound-based ambulatory in vitro fertilization program. Fertil Steril 1988, 49:468–78.

[35] Phelps JY, Levine AS, Hickman TN, Zacur HA, Wallach EE, Hinton EL: Day 4 estradiol levels predict pregnancy success in women undergoing controlled ovarian hyperstimulation for IVF. Fertil Steril 1998, 69:1015–9.

[36] Fisher S, Grin A, Paltoo A, Shapiro HM: Falling estradiol levels as a result of intentional reduction in gonadotrophin dose are not associated with poor IVF outcomes, whereas spontaneously falling estradiol levels result in low clinical pregnancy rates. Hum Reprod 2005, 20:84–8.

[37] Michalson KT, Groban L, Howard TD, Shively CA, Sophonsritsuk A, Appt SE, Cline JM, Clarkson TB, Carr JJ, Kitzman DW, Register TC: Estradiol Treatment Initiated Early After Ovariectomy Regulates Myocardial Gene Expression and Inhibits Diastolic Dysfunction in Female Cynomolgus Monkeys: Potential Roles for Calcium Homeostasis and Extracellular Matrix Remodeling. J Am Heart Assoc 2018, 7:e009769.

[38] Kahveci B, Melekoglu R, Evruke IC, Cetin C: The effect of advanced maternal age on perinatal outcomes in nulliparous singleton pregnancies. BMC Pregnancy Childbirth 2018, 18:343.

[39] Odibo AO, Nelson D, Stamilio DM, Sehdev HM, Macones GA: Advanced maternal age is an independent risk factor for intrauterine growth restriction. Am J Perinatol 2006, 23:325–8.

[40] Gaillard R, Bakker R, Steegers EA, Hofman A, Jaddoe VW: Maternal age during pregnancy is associated with third trimester blood pressure level: the generation R study. Am J Hypertens 2011, 24:1046–53.

[41] Matthiesen L, Berg G, Ernerudh J, Ekerfelt C, Jonsson Y, Sharma S: Immunology of preeclampsia. Chem Immunol Allergy 2005, 89:49–61.

[42] Lokki AI, Heikkinen-Eloranta JK, Laivuori H: The Immunogenetic Conundrum of Preeclampsia. Front Immunol 2018, 9:2630.

[43] Han X, Ghaemi MS, Ando K, Peterson LS, Ganio EA, Tsai AS, Gaudilliere DK, Stelzer IA, Einhaus J, Bertrand B, Stanley N, Culos A, Tanada A, Hedou J, Tsai ES, Fallahzadeh R, Wong RJ, Judy AE, Winn VD, Druzin ML, Blumenfeld YJ, Hlatky MA, Quaintance CC, Gibbs RS, Carvalho B, Shaw GM, Stevenson DK, Angst MS, Aghaeepour N, Gaudilliere B: Differential Dynamics of the Maternal Immune System in Healthy Pregnancy and Preeclampsia. Front Immunol 2019, 10:1305.

[44] Pitkin RM, Witte DL: Platelet and leukocyte counts in pregnancy. JAMA 1979, 242:2696–8.

[45] Canzoneri BJ, Lewis DF, Groome L, Wang Y: Increased neutrophil numbers account for leukocytosis in women with preeclampsia. Am J Perinatol 2009, 26:729–32.

[46] Brabin BJ: An analysis of malaria in pregnancy in Africa. Bull World Health Organ 1983, 61:1005–16.

[47] Archibald HM: The influence of malarial infection of the placenta on the incidence of prematurity. Bull World Health Organ 1956, 15:842–5.

[48] Walker PG, Griffin JT, Cairns M, Rogerson SJ, van Eijk AM, ter Kuile F, Ghani AC: A model of parity-dependent immunity to placental malaria. Nat Commun 2013, 4:1609.

[49] Ritzel RM, Patel AR, Spychala M, Verma R, Crapser J, Koellhoffer EC, Schrecengost A, Jellison ER, Zhu L, Venna VR, McCullough LD: Multiparity improves outcomes after cerebral ischemia in female mice despite features of increased metabovascular risk. Proc Natl Acad Sci U S A 2017, 114:E5673–E82.

[50] Ali AA, Rayis DA, Abdallah TM, Elbashir MI, Adam I: Severe anaemia is associated with a higher risk for preeclampsia and poor perinatal outcomes in Kassala hospital, eastern Sudan. BMC Res Notes 2011, 4:311.

[51] Hlimi T: Association of anemia, pre-eclampsia and eclampsia with seasonality: a realist systematic review. Health Place 2015, 31:180–92.

[52] Bilano VL, Ota E, Ganchimeg T, Mori R, Souza JP: Risk factors of pre-eclampsia/eclampsia and its adverse outcomes in low- and middle-income countries: a WHO secondary analysis. PLoS One 2014, 9:e91198.

[53] Lin L, Wei Y, Zhu W, Wang C, Su R, Feng H, Yang H, Gestational diabetes mellitus Prevalence Survey study G: Prevalence, risk factors and associated adverse pregnancy outcomes of anaemia in Chinese pregnant women: a multicentre retrospective study. BMC Pregnancy Childbirth 2018, 18:111.

[54] Obse N, Mossie A, Gobena T: Magnitude of anemia and associated risk factors among pregnant women attending antenatal care in Shalla Woreda, West Arsi Zone, Oromia Region, Ethiopia. Ethiop J Health Sci 2013, 23:165–73.

[55] Zimmermann MB, Hurrell RF: Nutritional iron deficiency. Lancet 2007, 370:511–20.

[56] Sato AP, Fujimori E, Szarfarc SC, Borges AL, Tsunechiro MA: Food consumption and iron intake of pregnant and reproductive aged women. Rev Lat Am Enfermagem 2010, 18:247–54.

[57] Abbaspour N, Hurrell R, Kelishadi R: Review on iron and its importance for human health. J Res Med Sci 2014, 19:164–74.

[58] Yang XP, Reckelhoff JF: Estrogen, hormonal replacement therapy and cardiovascular disease. Curr Opin Nephrol Hypertens 2011, 20:133–8.

[59] Marko KI, Simon JA: Clinical trials in menopause. Menopause 2018, 25:217–30.

[60] Borlaug BA, Redfield MM: Diastolic and systolic heart failure are distinct phenotypes within the heart failure spectrum. Circulation 2011, 123:2006–13; discussion 14.

[61] Upadhya B, Kitzman DW: Heart Failure with Preserved Ejection Fraction in Older Adults. Heart Fail Clin 2017, 13:485–502.

[62] Beale AL, Nanayakkara S, Segan L, Mariani JA, Maeder MT, van Empel V, Vizi D, Evans S, Lam CSP, Kaye DM: Sex Differences in Heart Failure With Preserved Ejection Fraction Pathophysiology: A Detailed Invasive Hemodynamic and Echocardiographic Analysis. JACC Heart Fail 2019, 7:239–49.

[63] Zeisler H, Jirecek S, Hohlagschwandtner M, Knofler M, Tempfer C, Livingston JC: Concentrations of estrogens in patients with preeclampsia. Wien Klin Wochenschr 2002, 114:458–61.

[64] Wan J, Hu Z, Zeng K, Yin Y, Zhao M, Chen M, Chen Q: The reduction in circulating levels of estrogen and progesterone in women with preeclampsia. Pregnancy Hypertens 2018, 11:18–25.

[65] Hertig A, Liere P, Chabbert-Buffet N, Fort J, Pianos A, Eychenne B, Cambourg A, Schumacher M, Berkane N, Lefevre G, Uzan S, Rondeau E, Rozenberg P, Rafestin-Oblin ME: Steroid profiling in preeclamptic women: evidence for aromatase deficiency. Am J Obstet Gynecol 2010, 203:477 e1–9.

[66] Jobe SO, Tyler CT, Magness RR: Aberrant synthesis, metabolism, and plasma accumulation of circulating estrogens and estrogen metabolites in preeclampsia implications for vascular dysfunction. Hypertension 2013, 61:480–7.

[67] Berkane N, Liere P, Oudinet JP, Hertig A, Lefevre G, Pluchino N, Schumacher M, Chabbert-Buffet N: From Pregnancy to Preeclampsia: A Key Role for Estrogens. Endocr Rev 2017, 38:123–44.

[68] Quinn MA, McCalla A, He B, Xu X, Cidlowski JA: Silencing of maternal hepatic glucocorticoid receptor is essential for normal fetal development in mice. Commun Biol 2019, 2:104.

[69] Quinn MA, Xu X, Ronfani M, Cidlowski JA: Estrogen Deficiency Promotes Hepatic Steatosis via a Glucocorticoid Receptor-Dependent Mechanism in Mice. Cell Rep 2018, 22:2690–701.

[70] Whirledge S, Cidlowski JA: Estradiol antagonism of glucocorticoid-induced GILZ expression in human uterine epithelial cells and murine uterus. Endocrinology 2013, 154:499–510.

[71] Whirledge S, Xu X, Cidlowski JA: Global gene expression analysis in human uterine epithelial cells defines new targets of glucocorticoid and estradiol antagonism. Biol Reprod 2013, 89:66.

[72] Wadhwa PD, Buss C, Entringer S, Swanson JM: Developmental origins of health and disease: brief history of the approach and current focus on epigenetic mechanisms. Semin Reprod Med 2009, 27:358–68.

[73] Rondo PH, Ferreira RF, Nogueira F, Ribeiro MC, Lobert H, Artes R: Maternal psychological stress and distress as predictors of low birth weight, prematurity and intrauterine growth retardation. Eur J Clin Nutr 2003, 57:266–72.

[74] Durousseau S, Chavez GF: Associations of intrauterine growth restriction among term infants and maternal pregnancy intendedness, initial happiness about being pregnant, and sense of control. Pediatrics 2003, 111:1171–5.

[75] Muhihi A, Sudfeld CR, Smith ER, Noor RA, Mshamu S, Briegleb C, Bakari M, Masanja H, Fawzi W, Chan GJ: Risk factors for small-for-gestational-age and preterm births among 19,269 Tanzanian newborns. BMC Pregnancy Childbirth 2016, 16:110.

[76] Huang YT, Lin HY, Wang CH, Su BH, Lin CC: Association of preterm birth and small for gestational age with metabolic outcomes in children and adolescents: A population-based cohort study from Taiwan. Pediatr Neonatol 2018, 59:147–53.

[77] Tong VT, England LJ, Rockhill KM, D’Angelo DV: Risks of Preterm Delivery and Small for Gestational Age Infants: Effects of Nondaily and Low-Intensity Daily Smoking During Pregnancy. Paediatr Perinat Epidemiol 2017, 31:144–8.

[78] Castrillio SM, Rankin KM, David RJ, Collins JW, Jr.: Small-for-gestational age and preterm birth across generations: a population-based study of Illinois births. Matern Child Health J 2014, 18:2456–64.

[79] Berghanel A, Heistermann M, Schulke O, Ostner J: Prenatal stress accelerates offspring growth to compensate for reduced maternal investment across mammals. Proc Natl Acad Sci U S A 2017, 114:E10658–E66.

